# The Rac1 homolog CED-10 is a component of the MES-1/SRC-1 pathway for asymmetric division of the *C. elegans* EMS blastomere

**DOI:** 10.1101/2024.04.04.588162

**Authors:** Helen Lamb, Małgorzata Liro, Krista Myles, McKenzi Fernholz, Holly Anderson, Lesilee S. Rose

## Abstract

Asymmetric cell division is essential for the creation of cell types with different identities and functions. The EMS blastomere of the four-cell *Caenorhabditis elegans* embryo undergoes an asymmetric division in response to partially redundant signaling pathways. One pathway involves a Wnt signal emanating from the neighboring P2 cell, while the other pathway is defined by the receptor-like MES-1 protein localized at the EMS/P2 cell contact, and the cytoplasmic kinase SRC-1. In response to these pathways, the EMS nuclear-centrosome complex rotates so that the spindle forms on the anterior-posterior axis; after division, the daughter cell contacting P2 becomes the endodermal precursor cell. Here we identify the Rac1 homolog, CED-10, as a new component of the MES-1/SRC-1 pathway. Loss of CED-10 affects both spindle positioning and endoderm specification. Although MES-1 is still present at the EMS/P2 contact in *ced-10* embryos, SRC-1 dependent phosphorylation is reduced. These and other results suggest that CED-10 acts downstream of MES-1 and upstream of, or at the level of, SRC-1 activity. In addition, we find that the branched actin regulator ARX-2 is enriched at the EMS/P2 cell contact site, in a CED-10 dependent manner. Loss of ARX-2 results in spindle positioning defects, suggesting that CED-10 acts through branched actin to promote the asymmetric division of the EMS cell.

## Introduction

Asymmetric cell division is the process by which dividing cells give rise to daughter cells with different identities and fates. This type of division occurs in all multicellular organisms and is necessary for cell type diversification during both embryonic development and adult tissue homeostasis (Inaba and Yamashita, 2012; Morin and Bellaiche, 2011; Venkei and Yamashita, 2018). A key aspect of asymmetric division is alignment of the mitotic spindle along a specific axis (Bergstralh and St Johnston, 2014; D’avino et al., 2005; di Pietro et al., 2016; Kotak, 2019; McNally, 2013). In many metazoan cell divisions requiring an oriented spindle, the microtubule pulling forces necessary to move the spindle are generated by the asymmetric localization of a conserved cortical complex. This force-generating complex contains the minus-end directed motor dynein, its partner dynactin, and the adaptor protein NuMA (LIN-5 in *C. elegans*, Mud in *Drosophila*). The complex can be recruited to the membrane or cortex by various adaptor and anchor proteins, and several different polarized cues have been identified that instruct spindle orientation (di Pietro et al., 2016; McNally, 2013; Morin and Bellaiche, 2011). For example, in many types of metazoan asymmetric divisions, the evolutionary conserved PAR proteins establish the polarity axis and influence the localization or activity of NuMA (LIN-5) and the force-generating complex. Wnt signaling pathways also act via NuMA (LIN-5) and dynein to align mitotic spindles with tissue polarity in several systems. However, for many cell types, the detailed mechanisms by which spindles are oriented remain to be elucidated (di Pietro et al., 2016; Gillies and Cabernard, 2011; Kotak, 2019).

The early *Caenorhabditis elegans* embryo serves as an excellent model for studying the molecular mechanisms of asymmetric division in different developmental contexts, as the embryo exhibits an invariant division pattern that includes multiple asymmetric divisions regulated by different cues (Griffin, 2015b; Pacquelet, 2017; Rose and Gonczy, 2014a). After fertilization, the zygote (or P0 cell) establishes molecular asymmetries that define the anterior and posterior embryonic poles. Following pronuclear meeting, the newly formed mitotic spindle rotates to align with the anterior-posterior axis and thus division creates an anterior somatic cell, AB, and a posterior germ-cell precursor, P1. The AB daughter divides symmetrically to give rise to ABa and ABp at the anterior and dorsal aspects of the embryo, respectively. The P1 cell divides asymmetrically to give rise to the EMS and P2 cells at the ventral and posterior aspects of the embryo, respectively. EMS and P2 both divide asymmetrically again (Fig. 1A)(Griffin, 2015a; Pacquelet, 2017; Rose and Gonczy, 2014b).

**Figure 1.**
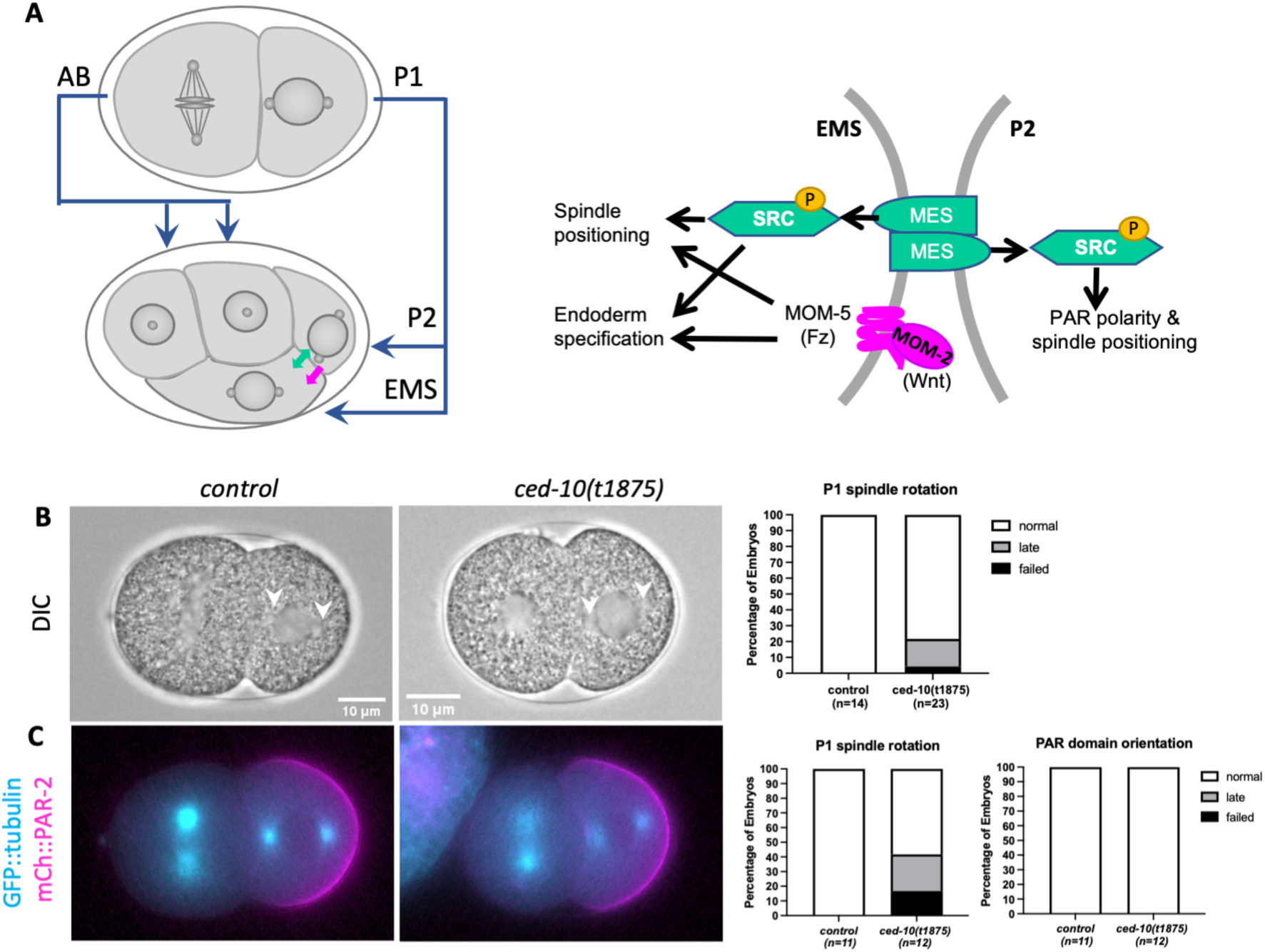
*ced-10* embryos undergo an asymmetric P1 division. (A) Diagram illustrating the progression from two cells to four cells in the early *C. elegans* embryo. In this and all images, anterior is to the left and posterior to the right, ventral is top and dorsal is bottom. The asymmetric division of the P1 cell results in differentially fated EMS and P2 daughters. The EMS/P2 cell contact is enlarged at right, with key components of the MES-1/SRC-1 (green) and Wnt (magenta) signaling pathways labeled. (B) Representative control GFP::tubulin and *ced-10(t1875)*; GFP::tubulin embryos imaged by DIC during the second division, illustrating normal nuclear centrosome complex orientation on the anterior-posterior axis in P1. Arrows point to centrosomes. Graph shows quantification of normal, late or failed P1 nuclear rotation (see Methods). (C) Representative epifluorescence images of mCh::PAR-2 domains and GFP::tubulin in control and *ced-10(t1875)* embryos at the time of P1 nuclear rotation. Graphs show quantification of P1 nuclear rotation in this background and PAR-2 domain orientation.

The asymmetric division of the P0 cell is instructed by PAR polarity. In these cells, PAR proteins antagonize one another’s cortical localization to form mutually exclusive anterior and posterior domains (Griffin, 2015b; Pacquelet, 2017; Rose and Gonczy, 2014a). In response to the formation of the PAR domains, a cytoplasmic gradient of cell fate determinants forms along the anterior-posterior axis. Downstream of PAR polarity cues, the force-generating complex described previously localizes asymmetrically on the cortex and exerts microtubule pulling forces that cause the nuclear-centrosome complex to rotate, such that the spindle forms along the axis of PAR asymmetry (Kotak, 2019; Pacquelet, 2017; Rose and Gonczy, 2014a). The resulting mitotic division gives rise to daughters that have inherited different quantities of PAR proteins and cytoplasmic fate determinants and therefore proceed to adopt different fates. In the asymmetric divisions of the posterior P1 daughter, mutually exclusive cortical PAR domains form and the nuclear-centrosome complex rotates onto the axis of PAR polarity. The mechanisms regulating nuclear rotation and unequal division of P1 are thought to be similar to those in the one-cell (Kotak, 2019; Pacquelet, 2017; Rose and Gonczy, 2014a).

In contrast, the asymmetry of the EMS division is generated by two partially redundant signaling pathways that require contact with EMS’s neighboring cell, P2: a well-described Wnt pathway and a less understood MES-1/SRC-1 pathway, involving the transmembrane tyrosine kinase-like receptor MES-1, the cytoplasmic tyrosine kinase SRC-1 (Fig. 1A; reviewed in (Maduro, 2017; Sawa, 2012)). In response to these pathways, the mitotic spindle orients on the anterior-posterior axis, which requires a rotation of the nuclear-centrosome complex from its initial left-right orientation (Fig. 1A). The anterior daughter of this division, MS, becomes a mesoderm progenitor, while the posterior daughter E, born adjacent to P2, becomes an endoderm progenitor.

Double mutant analysis has been the standard method for assigning genes to one of the partially redundant EMS asymmetric division pathways. Homozygous maternal loss of function for MES-1, SRC-1, or any of several Wnt pathway members causes a low rate of late or failed EMS spindle rotations (Bei et al., 2002; Rocheleau et al., 1997; Schlesinger et al., 1999; Thorpe et al., 1997; Walston et al., 2004). However, when a *mes-1* or *src-1* mutation is combined with mutation of a Wnt pathway component, the rate of spindle rotation failure increases to 90-100%.

In contrast, *mes-1;src-1* double mutants have the same low frequency of spindle rotation defects as *mes-1* and *src-1* single mutants. The MES-1 protein is localized to the EMS/P2 cell contact, apparently in both cells, as this localization requires cell-cell contact specifically between these cells (Bei et al., 2002; Berkowitz and Strome, 2000). Further, MES-1 is required for activation of SRC-1 at the EMS/P2 contact, based on the staining pattern of an antibody that detects SRC-1-dependent tyrosine phosphorylation (Bei et al., 2002). Thus, MES-1 and SRC-1 are part of a pathway that is functionally redundant with Wnt signaling in promoting EMS spindle positioning. The two pathways may converge on the conserved force generating complex described earlier. Loss of the dynein heavy chain (DHC-1), dynactin (DNC-1), or LIN-5 results in stronger spindle orientation defects than single mutants in either the MES-1/SRC-1 or Wnt pathways, suggesting that LIN-5 and dynein may act in both pathways (Liro and Rose, 2016; Zhang et al., 2007).

The induction of the E cell to form endoderm also depends on both the Mes and Wnt pathways. Endoderm fate specification requires the Wnt ligand (MOM-2 in *C. elegans*, expressed in the P2 cell), the transmembrane Frizzled receptor (MOM-5), and other conserved Wnt pathway components (Rocheleau et al., 1997; Schlesinger et al., 1999; Thorpe et al., 1997). Activation of this pathway ultimately results in nuclear export of the LEF-1/TCF-7 ortholog POP-1 in the E cell, which allows endoderm-specific gene expression in this cell (Maduro, 2017; Sawa, 2012). While mutation in either MES-1 or SRC-1 does not result in endoderm defects, mutation of either in combination with strong loss of function mutations in Wnt pathway components results in increased numbers of embryos without endoderm, and loss of SRC-1 was shown to enhance the POP-1 nuclear export defect of Wnt pathway mutants (Sugioka et al., 2011; Sugioka and Sawa, 2010; Sumiyoshi et al., 2011).

A previous study found that CED-10, one of three *C. elegans* Rac proteins, is required for spindle orientation in the ABar blastomere of the eight-cell embryo, which is a Wnt dependent pathway (Cabello et al., 2010). Rac proteins, a conserved subclass of the Rho GTPase family, are involved in cytoskeletal regulation across diverse developmental contexts and many organisms (Boreaux et al., 2007; Bustelo et al., 2007; Duquette and Lamache-Vane, 2014; Hall, 2012). The same *C. elegans* study reported that *ced-10* mutant embryos exhibited abnormal EMS spindle positioning in the four-cell embryo (Cabello et al., 2010). However, this phenotype was not characterized, nor was the relationship between CED-10 and the signaling pathways known to promote EMS spindle positioning. Therefore, we set out to define the genetic and molecular role CED-10 plays in this asymmetric division. Here we demonstrate that CED-10 works in parallel with the Wnt pathway in the EMS cell, where it acts downstream of MES-1 and upstream or at the level of SRC-1 for both spindle orientation and endoderm specification.

## Results

### The Rac1 homolog CED-10 is a member of the MES-1/SRC-1 pathway for EMS spindle positioning

Previous work showed that embryos from mothers homozygous for a null allele of *ced-10(t1875)*, hereafter referred to as *ced-10* mutant embryos, exhibited defects in the division of the ABar cell of the eight-cell embryo; this same study reported that CED-10 is required for spindle positioning in EMS, but no data were shown for this or earlier divisions (Cabello et al., 2010). We recently reported that *ced-10* mutant embryos exhibit normal nuclear and spindle positioning movements during the division of the one-cell embryo (Price et al., 2022). However, normal EMS division also requires the successful completion of the P1 division to produce properly fated P2 and EMS cells (Fig. 1A). Thus, here we first examined the P1 division in *ced-10* mutant embryos before proceeding to characterize the role of CED-10 in the EMS division. To facilitate scoring spindle positioning, we generated a strain with *ced-10(t1875)* in a GFP::tubulin background and compared embryos from this strain to control GFP::tubulin embryos.

In control embryos, the P1 nuclear-centrosome complex rotated onto the anterior-posterior axis prior to nuclear envelope breakdown (NEB) in all embryos (Fig. 1B). In comparison, 21% of *ced-10*; GFP::tubulin embryos had a late or failed P1 spindle rotation (n=23, Fig. 1B). PAR polarity is required for spindle orientation and for the identity of P1’s daughter cells, and so we next observed the localization of endogenously tagged mCherry::PAR-2 (mCh::PAR-2) in a *ced-10* strain also expressing GFP::tubulin. Late and failed P1 nuclear rotations were again seen for *ced-10* embryos in this background (Fig. 1C). Nonetheless, all embryos formed normally oriented PAR-2 domains, and the P2 cell inherited PAR-2 around its entirety in embryos with normal or late rotation (n=12, n=11 for control). A small proportion of *ced-10* embryos also inherited some PAR-2 in the EMS cell (1/12), but this was also seen in the control embryos (2/11). In *ced-10* embryos in which the P1 spindle oriented with normal or late timing, the EMS blastomere was larger than P2 and divided before P2, as in controls. Together these results suggest that CED-10 may have a subtle role in P1 nuclear rotation, but in the vast majority of embryos, the P1 cell divides asymmetrically.

We next examined the EMS division in *ced-10* embryos and observed that 43% of *ced-10* embryos had a late or failed EMS spindle rotation (Fig. 2A, B). CED-10 appears to act in a Wnt-dependent asymmetric division to orient the spindle of the ABar cell at the eight-cell stage (Cabello et al., 2010). We therefore scored spindle positioning defects in *ced-10* mutants depleted of MES-1, expecting that if CED-10 contributes to Wnt signaling for EMS spindle positioning, *ced-10; mes-1(RNAi)* embryos would show a higher rate of complete failures of EMS rotation. However, the proportion of failed and late rotations was not enhanced in *mes-1(RNAi);ced-10* embryos compared to the single mutants. Instead, the combination of RNAi depletion of MOM-2 (Wnt) and the *ced-10* mutant background increased the overall proportion of abnormal EMS spindle positions to 77% and increased the rate of failed EMS spindle rotation events to 53%, compared to 18% for *ced-10* alone (Fig. 2B). These results suggest that CED-10 plays a role in the MES-1/SRC-1 pathway for EMS spindle positioning.

**Figure 2.**
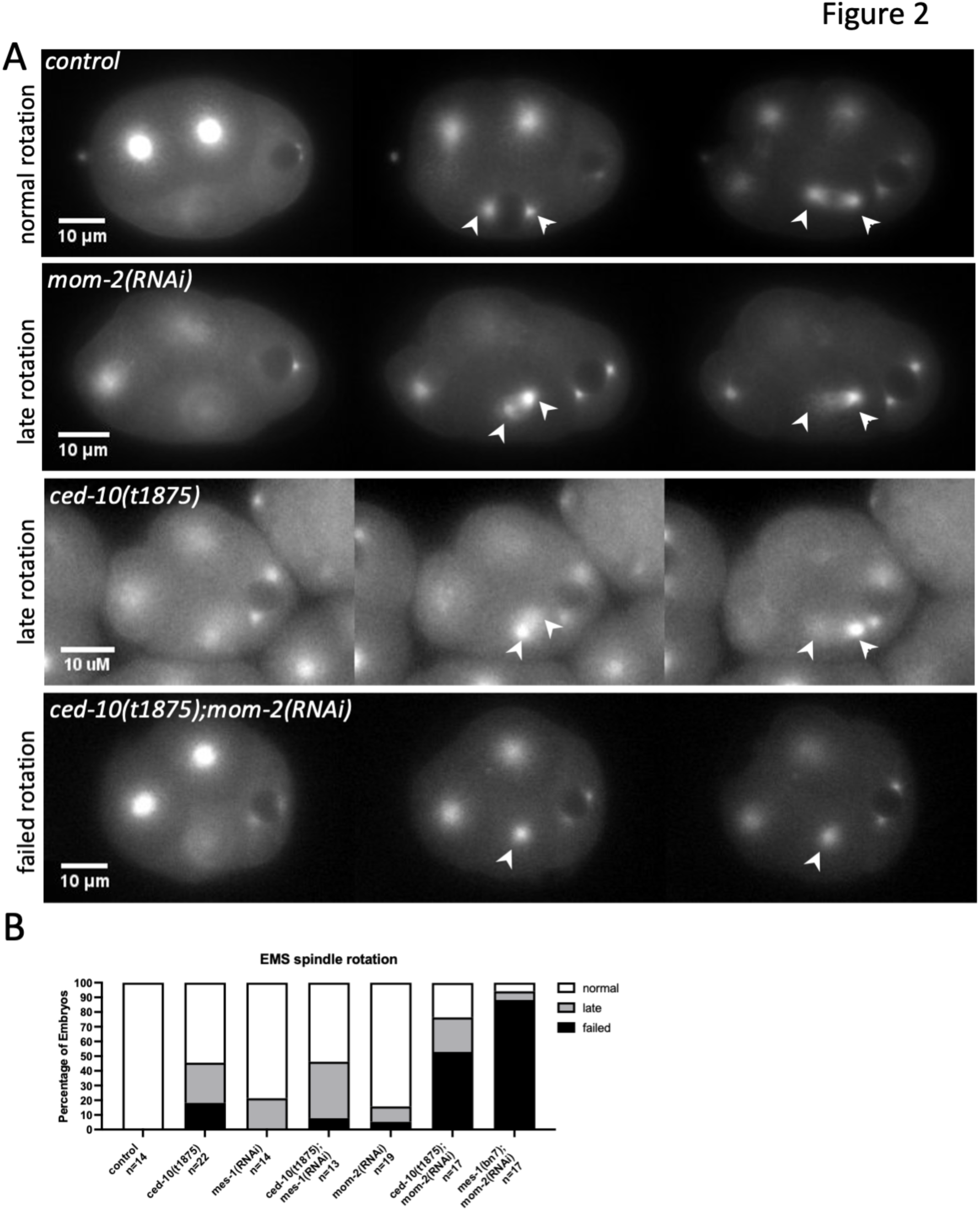
CED-10 acts in the MES-1/SRC-1 signaling pathway for EMS spindle positioning. (A) Representative stills from time-lapse images of GFP::tubulin embryos with the indicated genotypes, illustrating the three categories of EMS spindle orientation phenotypes observed. Arrowheads indicate EMS centrosomes. (B) Percentage of scored embryos for each indicated genotype with normal, late, or failed EMS rotation in a GFP::tubulin background. To ensure that the enhanced EMS defects were not indirectly caused by abnormal P1 division, only embryos with normal or late P1 spindle rotation were included in the analysis for *ced-10* and *ced-10; mom-2(RNAi)* embryos; 24% of *ced-10; mom-2(RNAi)* embryos had a late or failed P1 rotation, similar to that of *ced-10* alone.

### CED-10 is not required for spindle positioning in the P2 cell

The MES-1 protein is localized exclusively to the EMS/P2 cell contact in wild-type embryos (Berkowitz and Strome, 2000). The MES-1/SRC-1 signaling pathway, in addition to its importance in EMS spindle positioning and endoderm fate specification, is required for orienting PAR polarity and spindle positioning in the P2 cell (Fig. 1A)(Bei et al., 2002; Berkowitz and Strome, 2000). At birth, P2 inherits the posterior PARs such as PAR-2 uniformly around the cortex. Before division, a new PAR-2 domain forms, but in a different orientation to the previous P lineage divisions: The “posterior” PARs disappear from the dorsal side of the cell, including part of P2/ABp cell contact, such that they are present in a ventral “V” that includes the P2/EMS cell contact (Fig. 3); the “anterior” PARs localize to the reciprocal dorsal side of the cell (Arata et al., 2010; Rose and Gonczy, 2014a). The P2 spindle orients along this new axis of PAR asymmetry. In *mes-1(RNAi)* or mutant embryos, reciprocal PAR polarity domains form in the P2 cell but are misoriented in ∼70% of embryos; specifically, PAR-2 disappears from the P2/EMS as well as the P2/ABp cell contact in these embryos, resulting in a posterior PAR-2 domain. The P2 spindle is oriented along this anterior-posterior axis in ∼50% of *mes-1* mutant embryos, the proportion being lower because spindle and polarity are uncoupled in some embryos; many divisions are also equal in terms of daughter cell size (Arata et al., 2010).

**Figure 3.**
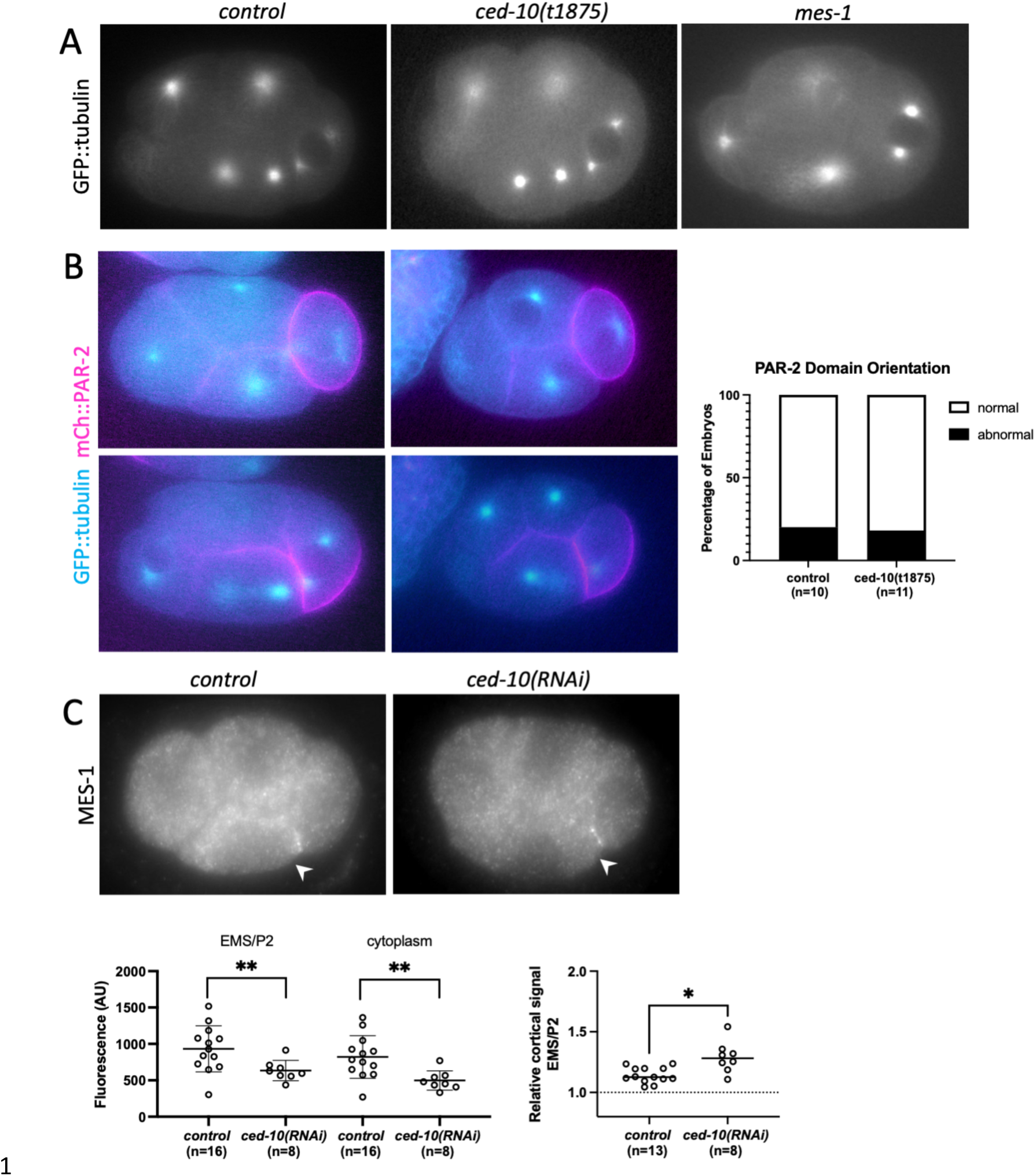
CED-10’s role in MES-1/SRC-1 signaling is downstream of MES-1. (A) Representative epifluorecence images of expressing GFP::tubulin embryos illustrating normal cueing of the P2 centrosome in control and *ced-10* embryos compared to *mes-1.* (B) Representative epifluorecence images of control and *ced-10* embryos expressing transgenic GFP::tubulin (cyan) and mCh::PAR-2 (magenta). Top panels show inheritance of PAR-2 around the entire periphery of P2; lower panels show formation of the PAR-2 domain. Graph shows percentage abnormal versus normal PAR-2 domains. (C) Epifluorescence microscopy of control and *ced-10(RNAi)* mNeonGreen::3xFLAG::MES-1 embryos immunostained for the FLAG epitope. Graphs below show quantification of absolute cortical and cytoplasmic intensities (left) and the cortical levels at the EMS/P2 contact relative to the cytoplasmic background (right).

If *ced-10* disrupts MES-1 function, then asymmetric division of the P2 cell should be affected. We first examined the P2 division from the same GFP::tubulin movies used above. Because the focal plane of these movies was optimized for scoring the EMS division, the final orientation or equalness in the P2 division was not always visible. Thus, we scored the initial orientation of the P2 nuclear-centrosome complex. The P2 nuclear-centrosome complex aligns with the PAR polarity axis through either centrosome migration or partial nuclear rotation where one centrosome becomes closely associated with P2/EMS cell contact (Arata et al., 2010; Berkowitz and Strome, 2000); the nuclear-centrosome complex and the subsequent spindle are thus oriented on a somewhat dorsal-ventral axis, approximately 45° relative to the anterior-posterior axis (Fig. 3A). For simplicity, we refer to the close association of the centrosome with the cell contact as “centrosome cueing”. We found that in control GFP::tubulin embryos, centrosome cueing occurred in all embryos, usually before EMS NEB (11/13; 2 embryos cued by EMS NEB + 40sec). Centrosome cueing also occurred in 100% of *ced-10(t1875)* embryos, within the same time range as controls (n=9) (Fig. 3A, B). In comparison, in the majority of *mes-1(bn7)* embryos, the P2 nuclear-centrosome complex did not cue to the cell contact (13/14), although in some embryos the centrosomes achieved a roughly dorsal-ventral orientation (Fig. 3A), as previously reported.

To further test for a *mes-1* like polarity phenotype, we examined P2 polarity in embryos expressing mCh::PAR-2 and GFP::tubulin. In the majority of control embryos and *ced-10(t1875)* mutant embryos, a PAR-2 domain formed on the ventral side of the cell as expected (Fig. 3B). In one case for each, PAR-2 remained uniform, and in one case for each, PAR-2 disappeared from the posterior of the P2 cell first. The latter is an opposite pattern to the defects noted above for *mes-1* embryos. Thus, *ced-10* does not affect the orientation of the PAR-2 domain in the P2 cell. In this data set, we again observed that in 100% of *ced-10* embryos (n = 10), the P2 nuclear-centrosome complex cued to the P2/EMS cell contact, within the same time range as controls cued (n =10). Together these data indicate that polarity and orientation of the P2 division are not affected by loss of CED-10.

We also examined the localization of MES-1. Although endogenously tagged mNeonGreen::3xFLAG::MES-1 is not visible in live embryos until the eight-cell stage (Heppert et al., 2018), we found that anti-FLAG antibody staining showed the expected MES-1 localization at the EMS/P2 cell contact. MES-1 was also localized at the EMS/P2 contact in *ced-10(RNAi*) embryos (Fig. 3C). The absolute intensity at the contact and in the cytoplasm were lower in *ced-10* embryos compared to controls and thus CED-10 depletion may affect overall MES-1 levels. Nonetheless, a higher cortical intensity of MES-1 at the EMS/P2 contact, relative to the cytoplasm, was still present (Fig. 3C). These results indicate that CED-10 is not required for MES-1 localization to the EMS/P2 contact per se. Together with the observations of a normal P2 division in *ced-10* embryos, these results suggest that CED-10 does not affect MES-1 activity and acts downstream of MES-1 in the pathway.

### CED-10 is required for normal levels of SRC-1 dependent phosphotyrosine staining

We next set out to determine whether CED-10 affects SRC-1. Staining of four-cell *C. elegans* embryos with the Y99 antibody, which recognizes phosphotyrosine, results in an enriched signal at the EMS/P2 contact compared to other contacts, which is SRC-1-dependent. Further, *mes-1* mutants show a loss of the enrichment, but not cell contact staining, which supports the model that MES-1 activates SRC-1 (Bei et al., 2002). However, whether SRC-1 protein itself is enriched at that EMS/P2 contact relative to other contacts in the four-cell embryo has not been tested. We therefore examined the localization of SRC-1 using a strain with SRC-1 endogenously tagged with GFP (Zhu et al., 2020a; Zhu et al., 2020b). SRC-1::GFP was cortically localized to all cell contacts at the four-cell stage, but was not enriched at the EMS/P2 contact compared to other cell contacts (Fig. 4A). These results together with prior findings suggest that the asymmetric localization of SRC-1 kinase activity is caused by asymmetric activation of SRC-1 at the EMS/P2 contact. The cortical localization of SRC-1::GFP was not altered in *ced-10(RNAi)* embryos (Fig. 4A).

**Figure 4.**
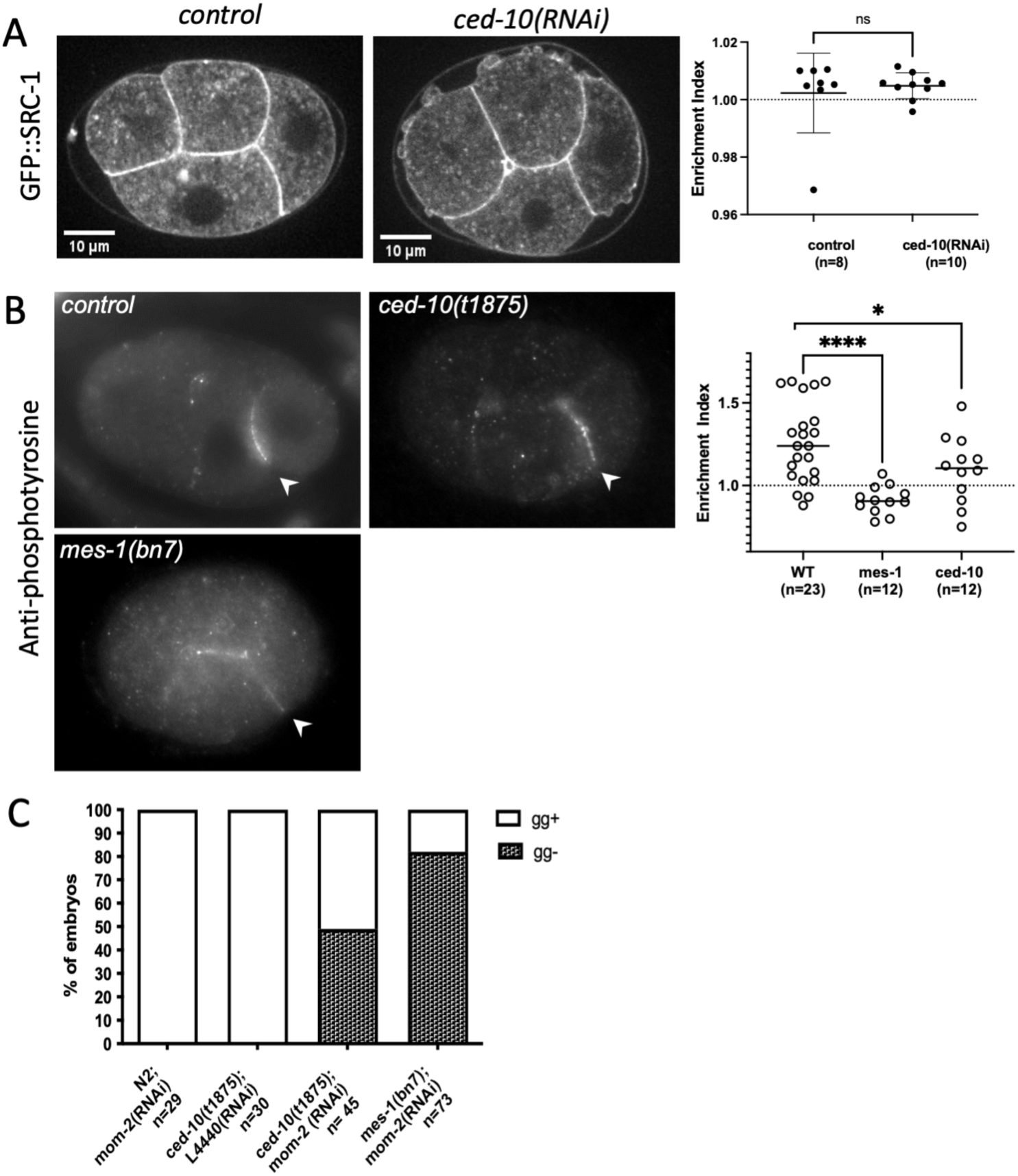
CED-10 is required for endoderm specification in parallel to Wnt signaling. (A) Representative confocal fluorescence images of GFP::SRC-1 in control and *ced-10(RNAi)* embryos. Graph shows the “enrichment index”, the level at the EMS/P2 contact compared to the level at the EMS/ABp contact. (B) Representative epifluorecence images of embryos stained for SRC-1-dependent phosphotyrosine in the indicated genotypes. Graph shows the “enrichment index” as for (A). (C) Proportion of embryos with and without gut granules in the indicated genotypes.

Four-cell embryos were also stained with the Y99 anti-phosphotyrosine antibody. While Y99 staining was still enriched at the EMS/P2 contact compared to the EMS/ABp contact in some *ced-10(t1875)* embryos, many embryos showed less enrichment than in wild-type (Fig. 4B). However, the average enrichment of the *ced-10* embryos was higher than for *mes-1* embryos, which had no enrichment of Y99 signal at the EMS/P2 contact, as previously reported (Bei et al., 2002).

In addition to EMS spindle positioning, MES-1 and SRC-1 are required for endoderm fate specification in the E daughter cell in parallel with Wnt signaling. That is, although loss of MES-1 or SRC-1 alone does not affect the formation of intestinal cells, loss of either enhances the variable intestine-minus phenotypes of Wnt pathway mutants (Bei et al. 2002). Since our results suggested that CED-10 affects SRC-1 activity at the EMS/P2 contact, we therefore asked whether loss of CED-10 affects the formation of intestinal tissue in late-stage embryos. As seen in prior studies (Bei et al., 2002; Rocheleau et al., 1997; Thorpe et al., 1997), we found that *mom-2(RNAi)* embryos did not hatch but did form intestinal tissue as scored by the presence of gut granules (Fig. 4C), while the majority of *mom-2(RNAi); mes-1(bn7)* embryos did not form intestinal tissue. All *ced-10(t1875)* embryos had intestinal tissue, but a large proportion of *ced-10(t1875)*; *mom-2(RNAi)* embryos did not exhibit gut granules (Fig. 4C). These data indicate that CED-10 is required for proper specification of endoderm tissue in parallel with Wnt signaling.

Together with the results of anti-phosphotyrosine staining, this observation suggests that CED-10 acts upstream of both EMS spindle positioning and endoderm fate specification by promoting enhanced SRC-1 activity or the accumulation of SRC targets at the EMS/P2 contact.

### CED-10 and ARX-2 are localized at cell contacts at the four-cell stage

To gain further insight into the function of CED-10 in MES-1/SRC-1 signaling, we characterized the localization of GFP::CED-10 in four-cell embryos, using an integrated GFP::CED-10 transgene under transcriptional control by the *ced-10* promoter (Ziel et al., 2009). This transgene was previously shown to rescue the defects of *ced-10 (n1993),* which is a viable allele of *ced-10* (Lundquist et al., 2001). We crossed this transgene into the *ced-10* null background and found that GFP::CED-10; *ced-10(t1875)* embryos appeared less round than *ced-10* mull embryos, and embryonic lethality was sufficiently rescued to maintain the strain homozygous for *ced-10*. In early embryos, GFP::CED-10 was present on all cell-cell contacts including at the four-cell stage (Fig. 5A). The cell contact signal and the cytoplasmic signal were both reduced to background levels by RNAi of *ced-10.* Although *GFP::CED-10; ced-10(t1875)* embryos exhibited many small cortical protrusions throughout the first few mitotic divisions, similar to *ced-10(t1875)* embryos (Price et al., 2022), *GFP::CED-10; ced-10(t1875)* embryos did not exhibit EMS division orientation defects. In addition, *GFP::CED-10; ced-10(t1875); mom-2(RNAi)* embryos did not show the higher rate of defective spindle orientations observed for *ced-10(t1875); mom-2(RNAi)* embryos examined in parallel (Fig. 5B). These results indicate that GFP::CED-10 is functional in the early embryo for EMS spindle positioning. Quantification of the relative cortical intensity of GFP::CED-10 at the EMS/P2 contact revealed a very slight enrichment compared to the EMS/ABp contact on average (Fig. 5C). The GFP::CED-10 localization pattern is consistent with a cortex-localized mechanism of action for CED-10 in EMS.

**Figure 5.**
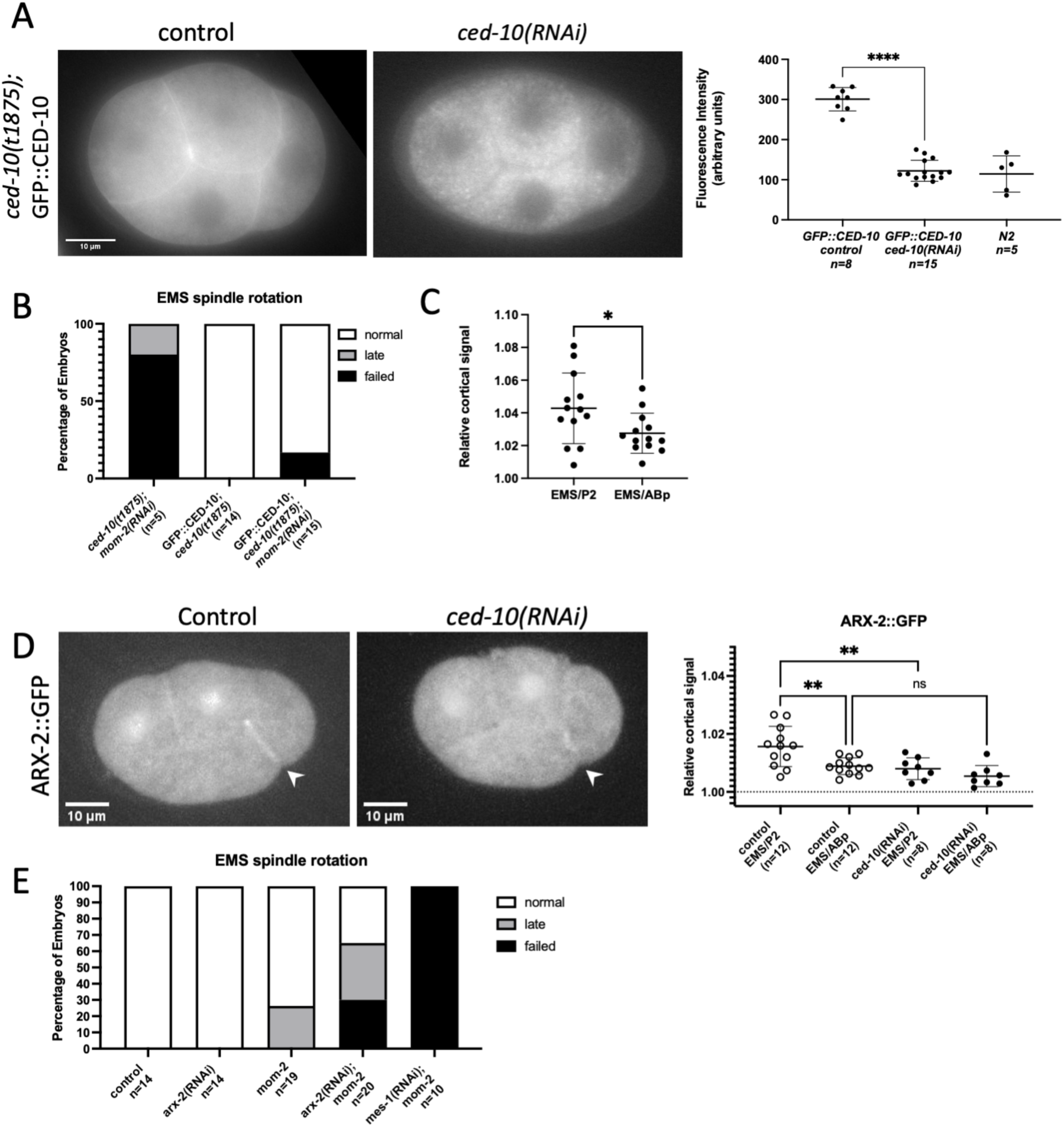
GFP::CED-10 and ARX-2::GFP are cortically localized in four-cell embryos. A) Representative wide-field fluorescence image of GFP::CED-10 in embryos treated with control RNAi (empty L4440 vector) or *ced-10* RNAi. Graph at right shows quantification of total background-corrected embryonic fluorescence intensity of GFP::CED-10 rescue embryos with and without *ced-10(RNAi)*. (B) Percentage of scored embryos with normal, late, and failed EMS spindle rotations for the indicated genotypes. (C) Quantification of relative cortical CED-10::GFP fluorescence intensity, normalized to cytoplasm, at both the EMS/P2 and EMS/ABp contacts. (D) Representative maximum Z projections from confocal fluorescence microscopy of ARX-2::GFP in control and *ced-10(RNAi)* embryos. Graph shows relative cortical ARX-2::GFP fluorescence intensity, normalized to cytoplasm, at both the EMS/P2 and EMS/ABp contacts in the indicated genotypes. (E) Percentage of scored embryos with normal, late, and failed EMS spindle rotations for the indicated genotypes.

Rac proteins can signal through the WAVE complex or WASp proteins, which both activate the Arp2/3 complex to nucleate branched actin (Saenz-Narciso et al., 2016; Shakir et al., 2008). Therefore, we hypothesized that CED-10 contributes to EMS asymmetric division by promoting branched actin at the EMS-P2 contact. Although prior studies have shown that branched actin reporters localize to the cell surface at the one-cell stage and to cell-cell contacts during ventral enclosure (Chan et al., 2018; Sawa et al., 2003), the localization of branched actin at the four-cell stage has not been reported. We visualized ARX-2, the Arp2 member of the Arp2/3 branched actin nucleating complex, with an integrated ARX-2::GFP fusion transgene under transcriptional control by the *arx-*2 promoter (Sarov et al., 2012). Consistent with our hypothesis, ARX-2::GFP was present at all cortical surfaces of the embryo and was also enriched at the EMS/P2 contact relative to the EMS/ABp contacts (Fig. 5D). Furthermore, the cortical localization of ARX-2::GFP at the EMS/P2 contact was decreased with *ced-10(RNAi)*, suggesting that CED-10 increases the levels of branched actin at the EMS/P2 cell-cell contact (Fig. 5D).

### ARX-2 is required for EMS spindle positioning

To test whether ARX-2 and thus branched actin contributes to EMS spindle positioning, we first depleted ARX-2 by RNAi in GFP::tubulin; *mom-*5*(zu193)* (*Frizzled* null) embryos, one of the standard Wnt mutant backgrounds used in studies of EMS spindle positioning (Cabello et al., 2010; Liro and Rose, 2016). Unexpectedly, we found that *mom-5(zu193); arx-2(RNAi)* embryos displayed a high rate of P1 spindle rotation defects, with 56% completely failed rotations, compared to 8% for *mom-5(zu193)* and 7% for *arx-2(RNAi)* GFP::tubulin embryos.

EMS spindle positioning was then scored in only the embryos with a normal or late rotation, which would allow for normal fating of the EMS and P2 blastomeres. In this subset, depletion of ARX-2 in a *mom-5(zu193)* background appeared to enhance the rate of defective spindle rotations observed in *mom-5* alone (Fig. S1); however, given the small sample size it is difficult to conclude from these data whether ARX-2 is required for EMS spindle positioning.

MOM-5/Frizzled was recently shown to affect pulling forces in the one-cell embryo, independent of its role in Wnt signaling (Sugioka et al., 2017). Thus, we repeated RNAi of ARX-2 in embryos from *mom-2(or42)* (Wnt) mothers. In these experiments, we did not observe P1 spindle rotation defects for either *mom-2* embryos or *mom-2; arx-2(RNAi)* embryos; however, ARX-2 RNAi enhanced the EMS spindle positioning defects of *mom-2* embryos (Fig. 5E).

Together these results show that ARX-2 is required for spindle positioning in the EMS cell in a pathway parallel to the Wnt pathway, consistent with ARX-2 and branched actin acting downstream of CED-10 to promote spindle positioning in the MES-1/SRC-1 pathway. Given the observation of P1 defects in *arx-2(RNAi); mom-5* embryos, as well as P1 defects in some *ced-10* embryos as noted earlier, we also examined *ced-10* RNAi in the *mom-5*; GFP::tubulin background. We observed a complete failure of P1 rotation in 40% of *ced-10(RNAi); mom-5* embryos, an increase compared to *mom-5* alone (Fig. S1). However, *ced-10(t1875); mom-2(RNAi)* did not exhibit an increased proportion of completely failed or late P1 spindle rotations (Fig.S1). These results suggest the P1 failure phenotype is due to an interaction between the Frizzled ortholog MOM-5 and the branched actin cytoskeleton.

## Discussion

In this study, we demonstrate that the *C. elegans* Rac protein CED-10 plays an important role in the asymmetric division of the EMS cell of the early embryo, which is regulated by partially redundant Wnt and MES-1/SRC-1 signaling pathways. Our data are consistent with CED-10 being a novel member of the MES-1/SRC-1 pathway. Our genetic and localization analyses suggest that CED-10 acts downstream of MES-1 and upstream or at the level of SRC-1. First, MES-1 was still present at the P2/EMS contact site, and *ced-10* embryos did not exhibit the defects in PAR-2 polarity and spindle orientation that are known to result from loss of MES-1.

Thus, although we cannot rule out a redundant role in the P2 division, CED-10 does not appear to be essential for MES-1 activity. At the same time, *ced-10* embryos did show reduced enrichment of SRC-dependent phospho-tyrosine signal at the EMS/P2 contact. The antibody used for this immunostaining experiment recognizes phosphorylated tyrosine and is a readout for both SRC-1 autophosphorylation and phosphorylation of its targets (Bei et al., 2002); thus loss of CED-10 could affect the activation of either SRC-1 or its targets. A GFP-tagged version of SRC-1 was present at cell-cell contacts, but was not enriched at the EMS/P2 contact, nor was SRC-1 localization altered by *ced-10(RNAi)*. These data argue against a role in the asymmetric recruitment of SRC-1 itself to the EMS/P2 cell contact. Instead, we propose that CED-10 promotes the cortical recruitment or retention of active SRC-1 or its targets at the EMS/P2 contact. A detailed analysis of this role will require future work uncovering direct targets of SRC-1 in EMS, since only one target in the early embryo has thus far been proposed (Sumiyoshi et al., 2011).

Our studies also reveal that ARX-2 and thus branched actin are required for the asymmetric division of the EMS cell. The defects in spindle positioning in *arx-2(RNAi)* embryos were very similar to those exhibited by *ced-10* embryos, and ARX-2::GFP was present at higher levels at the EMS/P2 cell contact compared to other contacts in a manner dependent on CED-10. Thus CED-10 appears to act through the generation of branched actin to promote spindle positioning in the EMS cell. Interestingly, these experiments also uncovered an apparent genetic interaction between branched actin and the Frizzled ortholog MOM-5 at the two-cell stage. In the absence of MOM-5, branched actin depletion with RNAi against *arx-2* or *ced-10* caused a high rate of failed P1 spindle rotations. Previous work implicated the Frizzled/MOM-5 protein and the adenomatous polyposis coli (APC) homolog APR-1 in the regulation of microtubule dynamics during mitotic spindle positioning of the one-cell *C. elegans* embryo. This mechanism is dependent on PAR polarity but independent of the MOM-2 ligand (Sugioka et al., 2017; Sugioka et al., 2011). As the P1 cell divides asymmetrically in a PAR dependent manner, it is possible that Frizzled and APR-1 participate in a similar mechanism of regulating microtubule dynamics for P1 spindle rotation as well. Based on the enhanced rate of P1 defects in *arx-2; mom-5* double mutants, branched actin is likely required in a parallel pathway for either spindle positioning or PAR polarity. It has recently been shown that PAR polarity in the P1 cell is established through redundant mechanisms, at least one of which involves the actomyosin skeleton (Koch and Rose, 2023; Ng et al., 2023).

Prior work characterized a role for Rac1/CED-10 in spindle positioning of the ABar cell of the eight-cell *C. elegans* embryos (Cabello et al., 2010). Wnt signaling instructs the mitotic spindle of ABar to set up at a perpendicular angle to the orientation of the three other AB cell spindles. This ABar spindle orientation pathway, like the Wnt-dependent pathway for EMS spindle positioning, is transcription-independent (Schlesinger et al., 1999; Walston et al., 2004). In the ABar cell, Wnt signaling appears to be the predominant mechanism because mutations in the Wnt components cause failure of spindle orientation with high penetrance. Further, overexpression of CED-10 resulted in slight rescue of Wnt mutant spindle orientation defects, and genetic analysis showed that CED-10 acts downstream of Wnt signaling for cell-corpse clearance in the later embryo and distal tip cell migration in the larvae (Cabello et al., 2010). In contrast, our analyses show that CED-10 is a new component of the MES-1/ SRC-1 pathway for both endoderm specification and spindle positioning in the EMS cell, rather than part of the Wnt pathway. SRC-1 has also been shown to contribute to ABar spindle positioning, but its precise role is unknown, and MES-1 does not appear to be present in the AB cells. Together these observations and our findings raise the possibility that SRC-1, CED-10 and ARX-2 work together in the ABar cell to promote spindle position, but downstream of Wnt signaling.

The data presented on the role of CED-10 in the EMS division are very similar to those reported previously for the PIG-1 kinase. PIG-1 is a PAR-1 related kinase that has partially redundant roles with PAR-1 in the one-cell embryo. During the EMS division however, genetic analyses placed PIG-1 in the MES-1/SRC-1 pathway while PAR-1 appears to act in the Wnt pathway. Loss of PIG-1 resulted in defects in both endoderm specification and spindle positioning, as seen here for loss of CED-10. Further studies are required to determine the mechanistic links among SRC-1, CED-10, ARX-2 and PIG-1 during this asymmetric division in *C. elegans*, and it will be interesting to learn if this mechanism is utilized by other cells in *C. elegans* or in other organisms.

## Data availability

Data necessary for confirming the conclusions of this study are present within the manuscript text or figures; this research did not generate large data sets but raw data and images are available upon request. Strains are available upon request from the corresponding author or the CGC.

## Acknowledgements

We thank members of the Rose and McNally labs for helpful discussions, and Daniel A. Starr, Bruce W. Draper, and Jennifer Heppert for experimental advice. We are grateful to Guangshuo Ou, Martha Soto, Karen Oegema, Francis McNally, and Erik Lundquist for strains and plasmids. Many strains were obtained from the *Caenorhabditis* Genetics Center, which is funded by the National Institutes of Health Office of Research Infrastructure Programs (P40 OD010440) for strains. The 3i Marianas spinning disk confocal and the Nikon laser scanning confocal used in this study were purchased using a National Institutes of Health Shared Instrumentation Grant [1S10RR024543-01]. We thank the MCB Light Microscopy Imaging Facility, which is a UC-Davis Campus Core Research Facility, for the use of this microscope.

This work was funded by awards from the National Institutes of Health Grant[R01GM68744] and the National Institute of Food and Agriculture [CA-D*-MCB-6239-H] to LSR. Support for HL and ML was also provided by the UC Davis BMCDB Graduate Group and a National Institutes of Health training grant to HL [T32 GM 007377].

## Materials and Methods

### Strains

Nematodes were maintained on Modified Youngren’s Only Bacto-Peptone (MYOB) agar plates, seeded with *E. coli* OP50 using standard procedures (Brenner, 1974; Church et al., 1995; Stiernagle, 2006). Temperature sensitive strains were maintained at 16°C and others were maintained at ambient temperature (22-24°C).

**Table 1.**
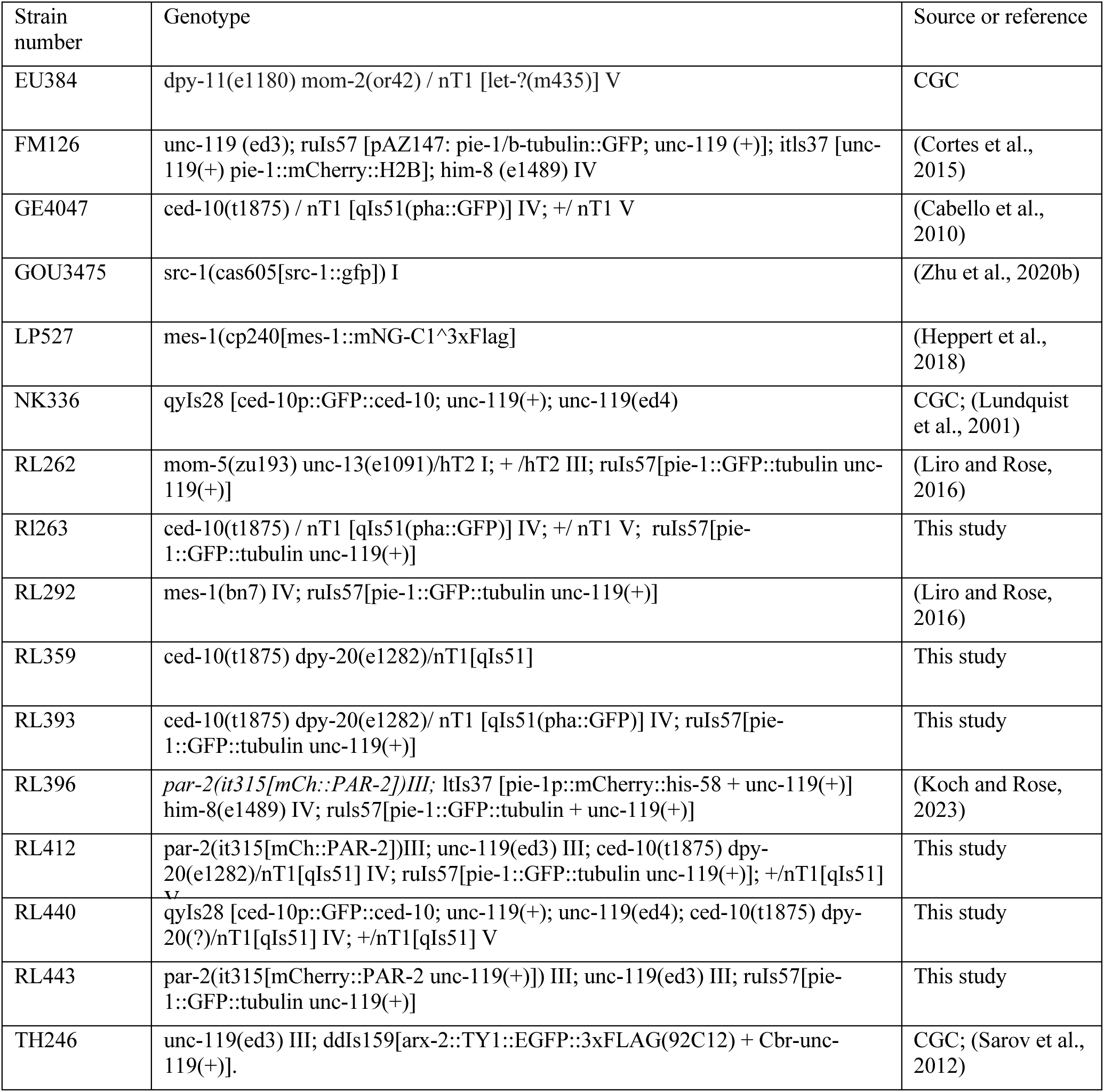
Strains used

### RNA interference

For all target genes, RNAi was performed by feeding as described (Ahringer, 2006), with the modification of adding IPTG to the bacterial culture, to a final concentration of 1mM, just prior to seeding the culture on the feeding plates. Plates were then kept in the dark at 4°C for no more than 5 days prior to use. The following RNAi feeding clones were obtained from the Ahringer library (Kamath et al. 2003): arx-2 (V-7M13), mom-2 (V-6A13), mes-1(X-5L23); the L4440 empty vector was used as control. For all conditions, worms were placed on plates seeded with RNAi bacteria at the large L3 through L4 stages and grown at 20°C unless otherwise noted, with imaging carried out at room temperature. For *arx-2*, worms were grown for 48-52hr; RNAi was judged to be effective if many small cortical extrusions were present in early embryos and there were unhatched siblings from same brood (Patel et al., 2008; Roh-Johnson and Goldstein, 2009). For *mom-2* RNAi in experiments to score spindle orientation, worms were grown for 36-48h and RNAi was considered effective if *mes-1(bn7);GFP::tubulin; mom-2RNAi* embryos filmed in parallel exhibited 80-100% abnormal spindle orientations; *mes-1* RNAi was carried out for 24-20hr, and considered *effective if mom-5(zu193); GFP tubulin; mes-1(RNAi)*embryos treated in parallel exhibited 80-100% abnormal spindle orientations as in (Liro and Rose, 2016). For *mom-2* and *mes-1* RNAi used to assess P granules, worms were grown at 16 for 48-72 hours, with RNAi effectiveness determined by enhancement of *mes-1(bn7)* or *mom-2(or42)* respectively as above.

The *ced-10* RNAi clone obtained from our library (IV-1J04) was found to contain no insert. An RNAi feeding construct was generated by subcloning the *ced-10* cDNA region of pPR37 (constructed by Peter Reddien, courtesy of Erik Lundquist) into the NcoI and SacI sites of the L4440 vector. The plasmid was transformed into HT115 and sequence confirmed. Defects in P1 and EMS spindle rotation in *mom-5(zu193); GFP::tubulin; ced-10(RNAi)* embryos confirmed that CED-10 knockdown with this plasmid was sufficient to induce an enhanced spindle phenotype, after 48-52h feeding at 20°C. Many small cortical extrusions were also present in these embryos, as reported for *ced-10(t1875)* (Price et al., 2022). In parallel experiments, knockdown efficacy was assessed by quantifying the depletion of fluorescent protein in GFP::CED-10; *ced-10(RNAi)* embryos after 48-52h at 20°C, which resulted in a reduction of the GFP signal to background N2 levels as described in the text; however, older embryos still showed CED-10::GFP. In addition, most *ced-10(RNAi)* embryos hatched, indicating that RNAi is not as strong as the null allele, but resembles published viable hypomorphic alleles (Kinchen et al., 2005; Lundquist et al., 2001). The following conditions were then used for RNAi of CED-10 for different experiments: For immunostaining for MES-1, 28-30hrs of feeding at room temperature, with the appearance of cortical blebs as a readout of RNAi efficacy. For GFP::SRC-1, ARX-2::GFP, and gut granule assays, feeding was carried out at 48-72 hours at 16C to allow for longer production of embryos. In addition to scoring for cortical blebs, enhancement of either the *mom-5* or *mom-2* spindle orientations defects was scored first as above, to ensure for RNAi effectiveness.

### Live epifluorecence microscopy

Embryos were removed from gravid hermaphrodites in 1x Egg Buffer, and mounted on 2% agarose pads under coverslips. For most experiments, embryos were observed on an Olympus BX60 microscope equipped with PlanApo N 60X, 1.42 NA oil immersion objective lens, a CoolLED light source, a Hamamatsu Orca 12-bit digital camera, and MicroManager software (Hardin, 2011; Strange et al., 2007).

To score P1, EMS, and P2 spindle positioning, images were acquired every 10 s with an exposure of 10 ms in bright field illumination. Focus was manually adjusted to follow centrosomes during each division. In strains expressing GFP::tubulin (all but *mom-2(or42)*), acquisition was switched to 488 nm light at approximately NEB of the ABa and ABp cells; this allows a more precise following of centrosomes and spindles in the EMS cell in mutants, because the centrosomes typically set up on the left-right axis in this cell. The same conditions were used to visualize GFP;;tubulin (10ms, 488nm) and PAR-2::mCherry (200ms, 560nm) localization, with the addition of fluorescence images at the start of the P1 cell cycle, at NEB/metaphase, and at P1 cytokinesis to score the P1 PAR-2 domain. To visualize GFP::CED-10, single-plane images were acquired with an exposure of 100 ms of 488nm light every 30 s.

For the *mom-2(or42)* single and double mutant experiments and gut granule experiments, embryos were prepared as above and imaged on an Olympus BX53 microscope equipped with PlanApo N 60X, 1.42 NA oil immersion lens, a Hamamatsu Orca Fusion BT camera, a SpectraX light engine, and motorized turret, all run by Olympus Cellsens software.

### Confocal microscopy

Embryos were removed from gravid hermaphrodites and mounted as above, and were observed using the spinning disc module of an Intelligent Imaging Innovations (3i) Marianas SDC Real-Time 3D Confocal-TIRF microscope fit with a Yokogawa spinning disc head, a 60x 1.4 numerical aperture oil-immersion objective, and EMCCD camera, at 100% 488 nm laser power. Acquisition was controlled by Slidebook 6 software (3i Incorporated). To visualize GFP::SRC-1, single-plane images were acquired with an exposure of 500 ms every 30 s. To visualize ARX-2::GFP, Z-stack images were acquired with a total depth of 14 um and a step size of 2 um at an exposure of 500 ms every 30 s.

### Scoring asymmetric division

Nuclear rotation was scored by following centrosome movements in brightfield and/or via tubulin fluorescence, with the angle of the nuclear-centrosome complex defined based on a line drawn through the two centrosomes relative to a line drawn on the anterior-posterior axis of the embryo. For the P1 cell, rotation was scored as “normal” if the nuclear-centrosome complex rotated to within 30° of the anterior-posterior axis (= 0°) prior to NEB, late if after NEB but prior to cytokinesis onset, and failed if rotation didn’t occur. With either normal or late rotation, one centrosome was also close to the AB/P1 cell contact after rotation. For the EMS cell, while in many cells the centrosomes are initially on the left-right axis occurs, we observed some cases of centrosome migration directly onto the anterior-posterior axis as previously reported (Liro and Rose, 2016). Thus, nuclear centrosome orientation was scored as normal is the centrosome were aligned with the anterior-posterior axis via either rotation or migration prior to NEB, and late if orientation occurred via spindle movements after NEB; in both cases the posterior centrosome was closer to the EMS/P2 contact at the end of an EMS division. A “failed” EMS rotation was defined as a spindle that was oriented more than 45° off the anterior-posterior axis in any direction at the end of division; the daughters cells in this case were in different focal planes after division. For the P2 cell, control GFP::tubulin embryos were filmed to determine the time at which close apposition of one centrosome to the EMS/P2 cell contact occurred, which can occur through direct centrosome migration or a slight rotation of the nucleus. The nuclear-centrosome complex and the subsequent spindle are thus oriented approximately 45° relative to the anterior-posterior axis, a slightly dorsal/ventral orientation. This “centrosome cueing” occurred in all control embryos, and the time ranged from 60sec before EMS NEB to 40 sec after in this data set. Centrosome position was then scored as normal if centrosome cueing occurred within this time range for GFP::tubulin strains.

Polarity was examined using mCh::PAR-2; GFP::tubulin expressing strains. For the P1 cell, embryos were scored as normal if a PAR-2 posterior domain was present by metaphase. Embryos in which PAR-2 was inherited abnormally from the first division (e.g. laterally in both AB and P1) were excluded from analysis, as the P1 cell would not be fated normally; this was observed in 1/11 control embryos. For the P2 cell, a “normal” PAR-2 domain was restricted to the ventral half to three-quarters of the cell; that is PAR-2 disappeared from at least the dorsal half of the ABp contact and the dorsal-most aspect of the outer posterior membrane of P2, and this occurred before P2 NEB. “Abnormal” P2 PAR-2 domains observed in both controls and *ced-10* embryos included uniform presence of mCherry::PAR-2 on the entire surface of P2 and the absence of mCherry::PAR-2 from the posterior cell surface only. P2 nuclear-centrosome orientation in PAR-2 strains was scored as above, but we noted more variability in the time of P2 centrosome cueing in this strain, with some control embryos cueing as late as EMS NEB + 160 sec, and thus *ced-10* embryos were scored as normal if they exhibited cueing within this time range.

To determine if embryos produced intestinal cells (an E cell derivative), embryos were mounted on agar pads as outlined for imaging of early divisions above. After filming some embryos to confirm spindle positioning phenotypes (e.g., in RNAi conditions), these and sibling embryos on agar pads were incubated in a moist chamber at 16°C for 18-24 hr, until control embryos had hatched. Polarization optics were then used to identify the presence of birefringent gut granules, which are a marker of intestinal differentiation (Laufer et al., 1980).

### Analysis of cortical fluorescence intensity from live imaging

Qualitative analysis of GFP::SRC-1 at the four-cell stage did not suggest any asymmetry at the EMS/P2 contact; quantifications of the relative cortical fluorescence of EMS/P2 contact compared to other contacts was measured from single images taken at the time of ABa/ABp NEB (scored by the loss of exclusion of fluorescent signal from the nucleus), as this is within the time period when published LIN-5 and Y99 asymmetries were visible. The “Segmented Line” tool in ImageJ was used to trace a 3-pixel-wide line along the entire EMS/P2 contact and other contacts, and the “Measure” function was used to obtain the mean fluorescence intensity of the the line. The same line traces were placed approximately 5 -10 pixels from the cell-cell contacts, avoiding the nucleus or centrosome, to sample mean cytoplasmic fluorescence intensity within the adjacent cells. The fluorescence intensity of each cell-cell contact was then divided by the mean cytoplasmic intensity of the two cells on either side of it to produce a relative cortical fluorescence intensity. The EMS/P2 “Enrichment Index” was calculated as the ratio of cytoplasm-normalized EMS/P2 contact fluorescence to the ratio of cytoplasm-normalized EMS/ABp contact fluorescence for each embryo. A value of 1 indicates no difference in relative cortical fluorescence intensity between the two cell-cell contacts, i.e., no enrichment.

For ARX-2::GFP, Maximum Z projections were generated for a timepoint 30-90 seconds prior to EMS NEB, as this appeared qualitatively to be the time with the strongest cell contact signal. Quantification of relative cortical intensity and enrichment were measured as above. For CED-10::GFP epifluorescence images, measurements of relative cortical intensity and enrichment were made on mid-focal plane images at ABa/ABp NEB as above, with the modification that a 2-pixel segmented line was used. For the examination of CED-10::GFP signal after RNAi of *ced-10*, the “freehand line” tool was used to trace around the embryo and the mean fluorescence measured.

### Immunostaining

Embryos were dissected in PBS from gravid hermaphrodites onto slides coated with poly-L-lysine, then they were freeze-cracked in liquid N2 according to (Duerr, 2006) To stain SRC-1 dependent phosphotyrosine as in (Bei et al., 2002), slides were immediately fixed in prechilled methanol at -20°C for 5 min, followed by 5 min in ice-cold acetone, air-dried for 5-10 min at room temperature, and then rehydrated in PBS for 5 min. Slides were incubated in 50 uL of blocking solution (5% w/v bovine serum albumin in PBS + 0.1% v/v Tween) in a moist chamber for 2 hr at room temperature. After wicking excess blocking solution away, samples were incubated in 50 uL of primary antibody solution (Santa Cruz Biotech mouse monoclonal PY99 diluted 1:200 in PBS + 1% w/v bovine serum albumin + 0.1% v/v Tween, overnight in a moist chamber at 4°C. Slides were washed 3x in PBS + 0.1% v/v Tween, then incubated in 50 uL secondary antibody solution (Alexa594-congugated goat monoclonal anti-mouse diluted 1:200 in PBS + 1% w/v bovine serum albumin + 0.1% v/v Tween) in a moist chamber for 2 hr at room temperature. After washing 3x in PBS + 0.1% v/v Tween, slides were incubated in DAPI/PBS followed by one wash in PBS; excess PBS was wicked away and samples were preserved with 20 uL of Vectashield mounting medium with a 20 x 40 mm coverslip sealed with nail polish. To stain mNG::3xFLAG::MES-1, the same protocol was used, omitting the acetone fixation and air-drying steps. The primary antibody was Invitrogen mouse monoclonal anti-FLAG, diluted 1:500. Images of samples were acquired in the focal plane of the EMS/P2 contact with an exposure of 200 ms of 488 nm light and 10ms of UV LIGHT on the Olympus BX60 microscope described above. Quantification of cortical intensity and enrichment was carried out in ImageJ as described above for SRC-1::GFP. For mNG::3xFLAG::MES-1 staining, a Maximum Z projection of two mid focal plane images from each four-cell stage embryo was used. For Y99 staining, quantification was carried on single-plane images from embryos in prophase through early anaphase, as determined by DAPI staining.

---

**Figure S1.**
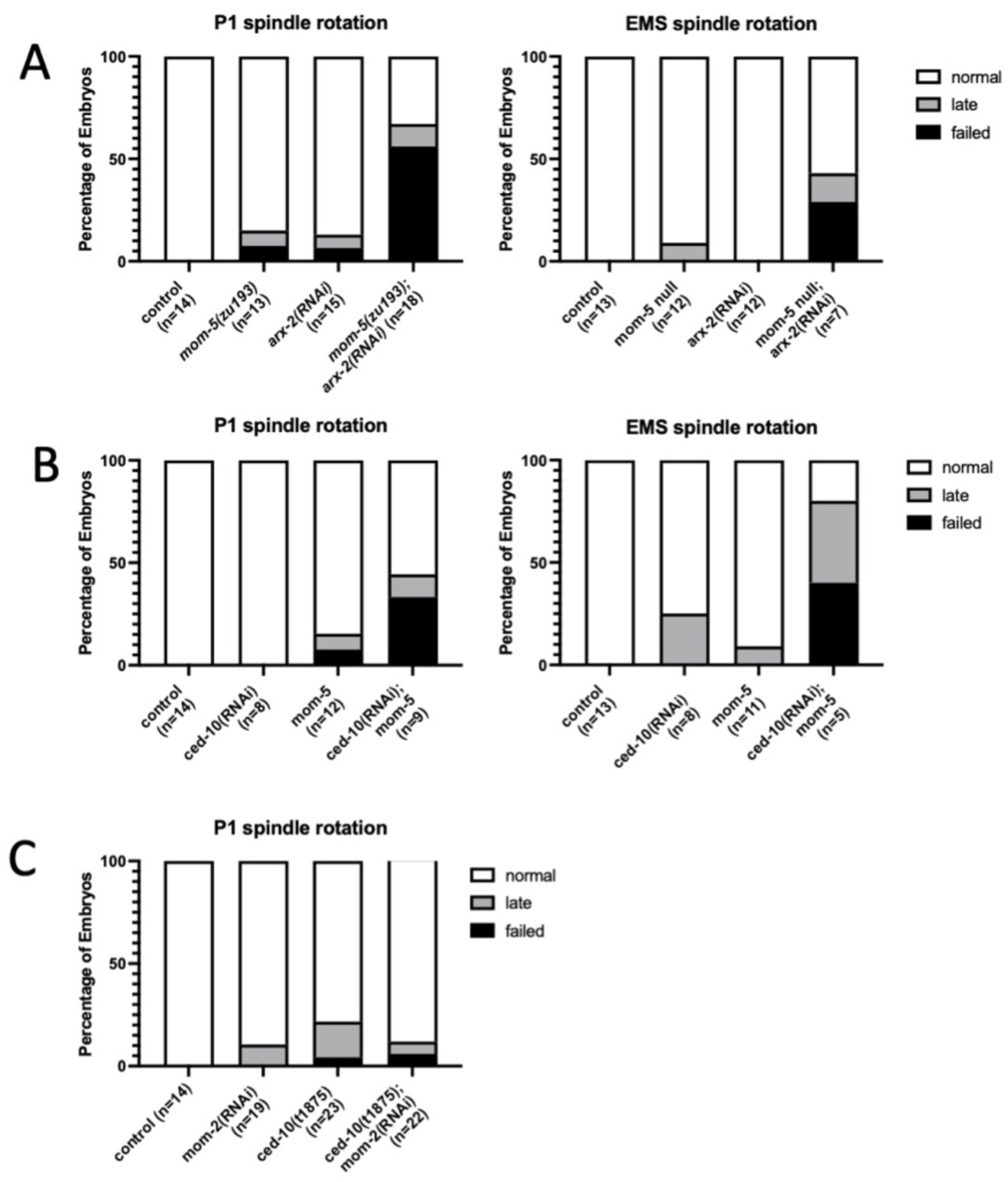
ARX-2 and CED-10 genetically interact with MOM-5(Frizzled) during P1 nuclear rotation. Percentage of scored embryos with normal, late, and failed EMS spindle rotations for the indicated genotypes. (A) Examination of P1 and EMS nuclear rotation in *arx-2; mom-5* double mutants. (B) P1 and EMS nuclear rotation in *ced-10; mom-5* double mutants. (C) P1 rotation in *ced-10; mom-2* double mutants.

